# An interaction between Yorkie and Taiman transcriptional coactivators regulates niche-to-stem cell BMP signaling in the female *Drosophila* germline

**DOI:** 10.64898/2026.10.28.762554

**Authors:** Chloe Wells, Victoria C. Placentra, Shilpi Verghese, Tod Butenschon, Kenneth H. Moberg

## Abstract

Communication between stem cells and their specialized niche microenvironments is vital for organism development and tissue homeostasis. However, mechanisms that limit intracellular signaling within a given niche and thus maintain stem cell fate are not fully understood. The Hippo and Ecdysone nuclear receptor (EcR) pathways are well conserved and individually required in niche cells for the establishment and maintenance of stem cell pools in *Drosophila*. These pathways are linked in the nucleus via a physical interaction between their respective transcriptional coactivators Yorkie (Yki) and Taiman (Tai). Within the female *Drosophila* germline niche, Yki and Tai each regulate behavior of stem cells and their progeny. It is not known whether these roles are mechanistically distinct or are a product of a physical interaction between Yki and Tai that enables a common niche-support role. Here we utilize a CRISPR mutant fly line containing a form of Tai unable to bind Yki (*tai^PPxA^*) to examine requirements for the Yki:Tai interaction *in vivo*. We find that the Yki:Tai interaction controls the GSC niche mainly through a role in escort cells (ECs), a key somatic cell-type that ensheathes the niche and regulates intercellular signaling. *tai^PPxA^* reduces EC cytoplasmic processes and allows the Dpp ligand to spread beyond its normally restricted range within the GSC niche, which elevates Dpp signaling and blocks differentiation of GSC progeny. In sum, these findings suggest that a Yki:Tai interaction facilitates crosstalk between the Hippo and EcR pathways within ECs and thus plays a critical role in maintaining intercellular signals that allow somatic cells to shape the niche microenvironment.

## Introduction

The unique ability of a fertilized egg to give rise to a complex multicellular organism capable of maintaining tissues throughout life and regenerating them after injury relies on stem cells. Stem cells undergo self-renewing divisions that maintain the stem cell population and generate daughter cells capable of replacing missing or damaged cells. Central to stem cell function are the specialized microenvironments they reside in, termed ‘niches’, that provide signaling cues and physical interactions to balance stem cell renewal and differentiation (rev. in Ohlstein et al., 2004). Genetic screens in tractable organisms have identified multiple signaling pathways that establish and maintain local niche environments (e.g., Hubbard and Schedl, 2019; Losick et al., 2011), but how these multiple inputs are integrated to drive the unique gene expression programs that sustain niches is not well defined.

One well-established system for studying niche-to-stem cell interactions is the *Drosophila melanogaster* female germline stem cell (GSC) niche located at the anterior tip of each ovariole in a structure termed the germarium (rev. in Nystul and Spradling, 2010). Each germarium houses two stem cell populations: germline stem cells (GSCs), which generate oocytes, and follicle stem cells (FSCs), which generate differentiated epithelial cells that encapsulate developing egg chambers. The somatic cells that constitute the GSC niche - including terminal filament (TF) cells, cap cells, and ensheathing escort cells (ECs) - regulate stem cell fate by modulating intercellular signaling through multiple pathways, including the Bone Morphogenetic Protein (BMP), Hippo, and Ecdysone (Ec) pathways (Konig et al., 2011; Xie and Spradling, 1998b). However, coordination and crosstalk between these pathways across multiple cell types in the germarium remains incompletely understood.

While some regulatory pathways intersect at the level of shared targets or through post-translational modification of key components, the Hippo and Ecdysone pathways are linked directly through a physical interaction between their respective transcriptional coactivator proteins Yorkie (Yki) and Taiman (Tai) (Wang et al., 2016; Zhang et al., 2015a). Yki is recruited to DNA by its obligate sequence specific DNA-binding partner Scalloped (Sd) (Goulev et al., 2008; Wu et al., 2008; Zhang et al., 2008; Zhao et al., 2008), while Tai is normally recruited to DNA by a complex of the Ec receptor (EcR) and its partner, the RXRα homolog Usp (Bai et al., 2000b; Yao et al., 1992). Tai can interact simultaneously with Yki and EcR through distinct interaction motifs (Zhang et al., 2015a). The EcR:Tai interaction requires a cluster of LxxLL motifs located in the central region of the Tai protein (Bai et al., 2000a), while the Yki:Tai interaction requires a pair of PPxY (proline-proline-x-tyrosine) motifs within the Tai C-terminal transactivation domain that bind two WW (tryptophan-tryptophan) domains present in Yki. Overgrowth of larval discs caused by a Yki transgene is enhanced by co-expression of wildtype *tai* (*UAS-tai^wt^*) and repressed by co-expression of a *tai* allele with Y-to-A edits in its PPxY motifs (*UAS-tai^PPxA^*) (Zhang et al., 2015a). Human homologs of Tai (NCOA1-3) associate with the mammalian Yki homolog, YAP, and as well as steroid hormone receptors such as the estrogen receptor (ER) (Kim et al., 2017; Kushner et al., 2020; Yuan et al., 2023; Zhu et al., 2019). Tai substitutes for NCOA1-3 in supporting testosterone-stimulated Androgen Receptor (AR) transcriptional activity in fly cells (Takeyama et al., 2004). Intriguingly, co-overexpression of Tai and Yki in wing epithelial cells upregulates mRNAs that differ from traditional Hippo and EcR target genes in this cell type and are instead associated with epithelial regeneration (e.g., *dIlp8*, Boone et al., 2016; Colombani et al., 2012; Garelli et al., 2012) or stem cell maintenance (e.g., *piwi*, Cox et al., 2000; Fagegaltier et al., 2016; Tang et al., 2023).

Independent studies of Yki and Tai roles in the female germline have determined that both factors are required to limit the range of BMP signaling within the germarium through roles in supporting escort cells (ECs) (Huang et al., 2017; Konig et al., 2011). The BMP ligands Decapentaplegic (Dpp) and Glass-bottom boat (Gbb) are normally produced only by cap cells and anterior ECs (also termed inner germarial sheath cells; IGCs), thus limiting activation of BMP signaling to proximal GSCs and thereby maintaining their stem cell identity (Chen and McKearin, 2003; Song et al., 2004; Xie and Spradling, 1998a). Posterior displacement of GSC daughters (also referred to as cystoblasts; CBs) moves them out of range of the BMP source, permitting CBs to express the pro-differentiation factor *bag-of-marbles* (*bam*) (Chen and McKearin, 2003; Song et al., 2004). Targeted depletion of Yki elevates production of BMP ligands, primarily Dpp, within ECs and reduces the size of EC cytoplasmic processes that normally extend into the germarium and insulate CBs from cap cell-derived BMP ligands (Huang et al., 2017). Notably, EC-specific depletion of either the obligate Yki-binding transcription factor Scalloped or the Yki-inhibitory kinases Hippo and Warts does not significantly alter CB differentiation (Huang et al., 2017), implying that Yki is regulated by non-canonical mechanisms in ECs. Parallel analysis of *tai* alleles indicates that Tai is also required in posterior ECs to limit activation of the BMP pathway to the anterior niche (König et al., 2011) and to maintain EC cytoplasmic processes that project into the germarium interior (Konig et al., 2011).

Here, we use a CRISPR-edited genomic *tai^PPxA^* allele to investigate whether similarities between the effect of *yki* and *tai* alleles on the female GSC niche simply reflect independent but overlapping functions of Tai and Yki proteins or alternatively reflect a Yki:Tai complex that acts as a single functional unit required to maintain GSC niche homeostasis. Our analysis of *tai^PPxA^* germaria reveals that Yki and Tai must be capable of associating with each other in order to support CB differentiation and limit expansion of the Dpp gradient. Comparison of the effect of *tai^wt^* and *tai^PPxA^* expression in different germarial cell types indicates that the Yki:Tai interaction is most significant in anterior ECs (i.e., IGCs). In sum, these data indicate that maintenance of the female *Drosophila* GSC niche relies on functional cooperativity between the Hippo and EcR transcriptional co-activator proteins Yki and Tai.

## RESULTS

### Tai requires its Yki-interaction motifs to support oogenesis

The Yki:Tai interaction involves two tryptophan-tryptophan (WW) domains in Yki and two proline-proline-x-tyrosine (PPxY) motifs in Tai (Zhang et al., 2015a). AlphaFold modeling predicts a high-confidence structure in which the Yki WW domains and Tai PPxY motifs are each located at the end of shortα-helical regions (i.e. W-helical-W and PPxY-helical-PPxY) which align in a crosswise parallel orientation that positions each PPxY motif opposite a WW domain (**Fig. 1a**). The first PPxY:WW interaction involves residues 1432-1435 (PPAY) in Tai and residues 321-338 (WW2) in Yki. The second PPxY:WW interaction involves residues 1476-1479 (PPMY) in Tai, and residues 252-269 (WW1) in Yki. Angstrom (Å) distances between α-carbon atoms (Ca-Ca) in both interaction pairs are consistent with a high-confidence AlphaFold model (**Fig. 1b**), and reveal a consistent alignment of amino acids between both WW:PPxY motifs (red boxes in **Fig. 1c**). The Proline^0^ of each Tai PPxY motif (position <u>P^0^</u>PxY) contacts multiple residues in the opposing WW domain, including the first tryptophan W^0^, Tyr^−11^, Thr ^−2^, and a polar residue (Ser/Gln) at position ^−^1. The PPxY P<u>^+1^</u> residue engages two threonine residues, Thr^−17^ and Thr^−2^. The variable PPxY residues (i.e., Ala^+2^ and Met^+2^) interact with polar residues (Ser/Thr) at positions ^−^2 and ^−^3 relative to W^0^. The PPxY Y^+3^ at position ^+^3 also contacts these Ser/Thr^−2^ polar residues, in addition to a basic residue (Arg/Lys) at position ^−^4 and a hydrophobic residue (Leu/Ile) at −9 (all relative to W^0^). Notably, these regions of Tai lie in a C-terminal transactivation domain that is distinct from the N-terminal PAS domain that binds Abrupt and the three centrally located LxxLL motifs that bind EcR (Bai et al., 2000a; Jang et al., 2009). Together with our prior interaction studies and IP/MS-MS analysis, which identified Yki as the only WW domain-containing protein among proteins that copurify with Tai (Zhang et al., 2015a), these AlphaFold data are consistent with a robust and specific interaction between Yki and Tai mediated by their two PPxY:WW interfaces.

**Figure 1.**
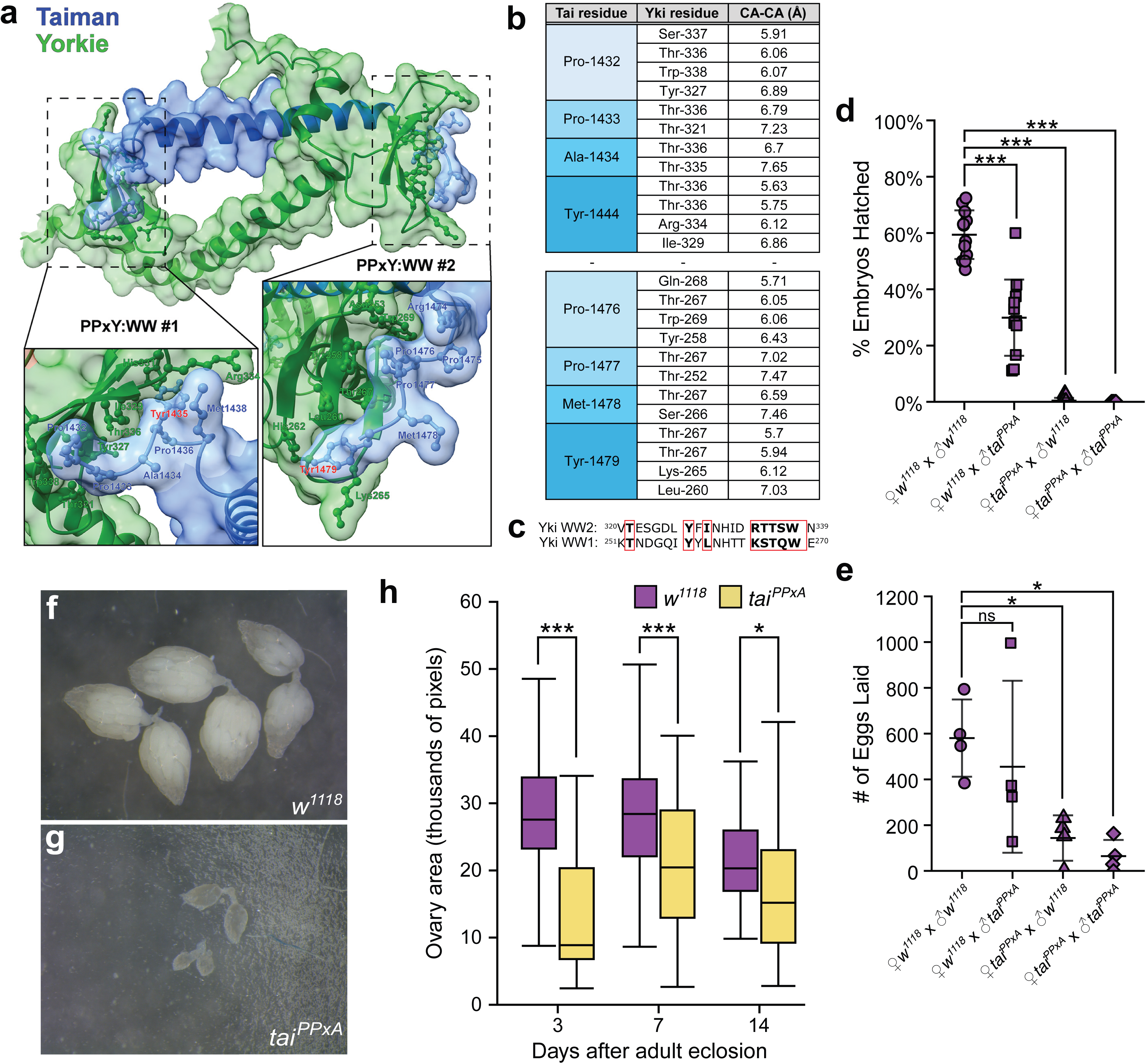
Tai requires its Yki-interaction motifs to support oogenesis. (**a)** Alphafold structure of the Taiman PPxY domain (blue) and the Yki WW domain (green). Note the intervening α-helices that are slightly rotated relative to each other. Dotted boxes indicate each PPxY:WW pair, which are magnified and rotated toward the viewer in boxes below. Key residues are labelled, including Try1435 and Tyr1479 that are edited to Ala in the *tai^PPxA^* allele. **(b)** Angstrom (Å) distances betweenα-carbon atoms in each PPxY:WW interaction pair. **(c)** Alignment of the Yki WW domains, with residues that make close contact with Tai boxed in red. **(d)** Egg counts from the indicated pairwise matings of control (*w^1118^*) and *tai^PPxA^* adults. **(e)** Embryo hatch rates of the indicated pairwise matings of control (*w^1118^*) and *tai^PPxA^*adults. **(f,g**) Representative images of control (*w^1118^*) and *tai^PPxA^* ovaries from unmated females 7 days after adult eclosion (AAE). **(h)** Quantification of ovary area (units of 10^3^ pixels) for control and *tai^PPxA^* females at 3, 7, and 14-days AAE. Sample sizes: control: n = 161, 116, 132; *tai^PPxA^*: n = 36, 40, 62. Statistical significance was assessed using Welch’s unpaired t-tests with Bonferroni correction. Error bars in (**d-e)** represent SEM. Boxes in (**f**) contain 50% of data points, whiskers contain upper and lower 25%. * = P ≤ 0.05, ** = P ≤ 0.005, *** = P ≤ 0.0005; ns, not significant.

To investigate the importance of the Yki:Tai interaction *in vivo*, we used CRIPSR/Cas9-mediated homology directed repair (HDR) to create a *tai* allele containing tyrosine-to-alanine substitutions in the two Tai PPxY motifs (PPAY^1435^-to-A and PPMY^1479^-to-A), hereafter referred to as *tai^PPxA^*. Following identification of successful repair events using an HDR template containing a *3x-P3>GFP* selection cassette, we subjected these chromosomes to multiple backcrosses and excised the *3x-P3>GFP* using a *heat-shock>Cre* line (BDSC #766). This yielded viable *tai^PPxA^* homozygote adults that are slightly reduced in size (data not shown), exhibit shortened lifespan (**Fig. S1a**) and reduced female fecundity (**Fig. 1d-e**). We chose to focus on the latter phenotype due to its severity and high penetrance. Female fecundity was quantified by counting the number of eggs laid by cohorts of mated females (**Fig. 1d**) and the fraction of eggs that hatched to produce L1 larvae in matings of *tai^PPxA^* and control (*w^1118^*) adults (**Fig. 1e**). *tai^PPxA^* females mated to control males lay fewer eggs than females in control crosses (p=0.043), and even fewer eggs when mated to *tai^PPxA^*males (p=0.029). In both mating schemes, very few eggs laid by *tai^PPxA^* females hatched as L1 larvae (respectively, p=3.7e-10 and p=4.9e-10) (**Fig 1e**). These data suggest that *tai^PPxA^* females produce fewer eggs and with reduced viability. The further reduction in egg production when *tai^PPxA^* females are mated to *tai^PPxA^* males indicates an additional *tai^PPxA^* paternal effect that was not investigated in this study.

To assess the effect of *tai^PPxA^* on the female reproductive system, we compared the structure of ovaries collected from unmated adult females (**Fig. 1f-h**) so as to analyze baseline GSC production and differentiation independent of the stimulatory effect of male sex-peptide (SP) on ecdysone-induced GSC proliferation (Ameku and Niwa, 2016b; Jung et al., 2025). Ovaries from *tai^PPxA^* females are dramatically decreased in size and have far fewer mature egg chambers (**Fig. 1f-g**). Given that mature egg chambers contribute substantially to ovarian volume, their absence likely explains the significantly reduced ovary area observed in *tai^PPxA^* mutants at 3, 7, and 14 days after adult eclosion (AAE) (p=1.2e-7, p=4e-4, p=0.01 respectively) (**Fig. 1h**). In sum, these data indicate that disrupting the Yki:Tai interaction decreases the number of mature eggs able to be produced and laid by *tai^PPxA^* mutant females.

### The Yki:Tai interaction is required for proper GSC differentiation, but not BC migration

The previously described roles of Yki and Tai in the non-cell-autonomous regulation of GSC differentiation (Huang et al., 2017; König et al., 2011) led us to examine the germarial landscape in *tai^PPxA^* mutants as a possible cause of the oogenesis defect. As illustrated in **Fig. 2a**, the germarium contains 2-3 GSCs, which contact cap cells that secrete factors that maintain stem cell identity, such as the BMP ligands Glass bottom boat (Gbb) and Decapentaplegic (Dpp). Moving posteriorly, ensheathing escort cells (ECs) extend cytoplasmic processes into the germarium interior that wrap around differentiating cystoblasts (CBs) and maintain their differentiation state (Decotto and Spradling, 2005; Schulz et al., 2002). Within the germarium, the GSC polarity marker α-spectrin (αSpec) protein coalesces in ‘dot spectrosomes’ that mark GSCs and their immediate cystoblast progeny in the anterior germarium, and ‘branched fusomes’ in further differentiated cysts in the mid-germarium (McKearin, 1997; Xie and Spradling, 1998a). Quantifying dot spectrosome number and fusome morphology (red in **Fig. 2a** cartoon) is a well-validated approach to assess the effects of mutations on GSC differentiation (rev. in Cuevas and Spradling, 1998).

**Figure 2.**
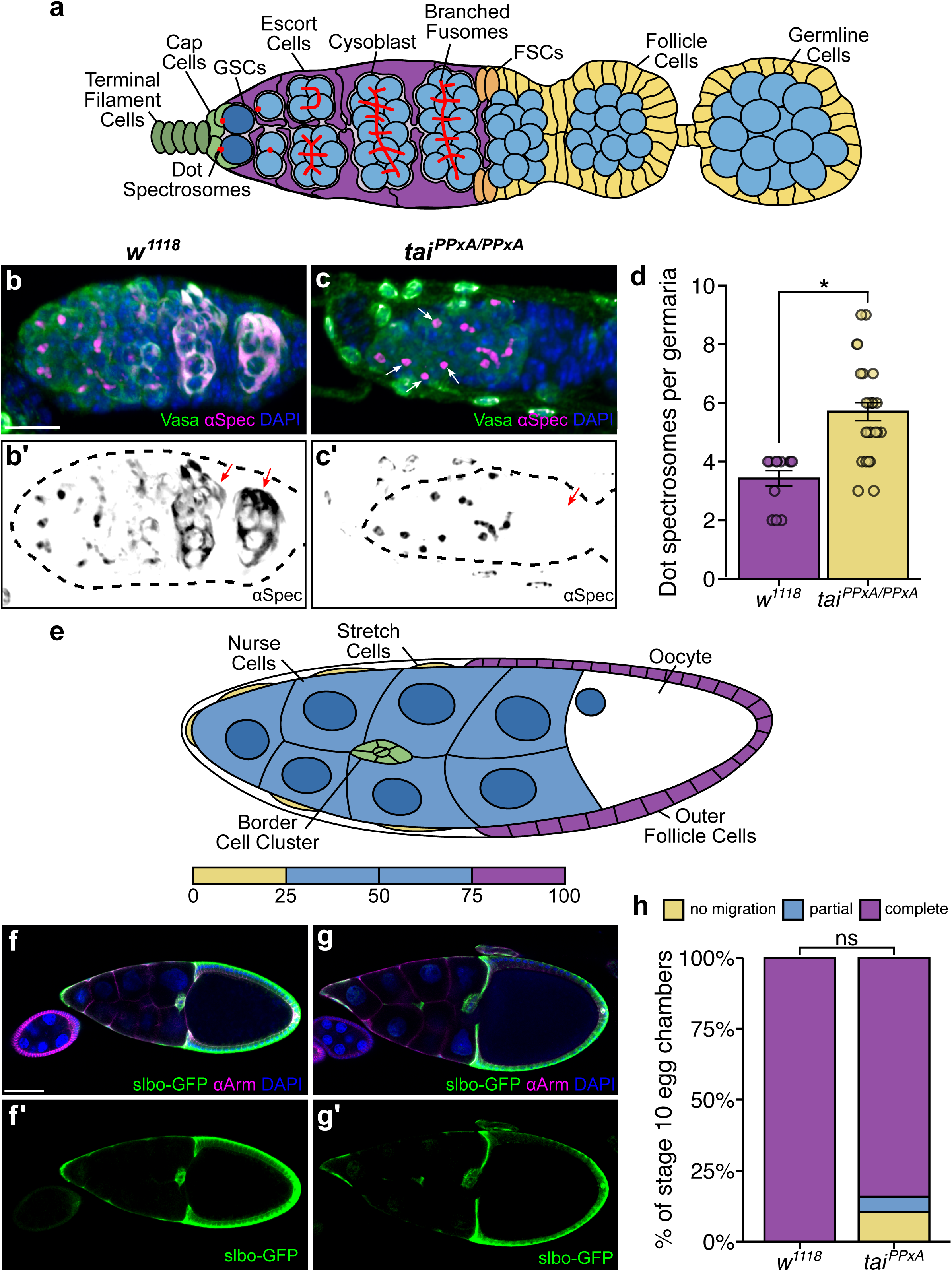
The Yki:Tai interaction is required for GSC differentiation, but not BC migration. **(a)** Diagram of the GSC niche (far left) and early phases of oocyte differentiation, with germaria regions I, IIa, IIb, and III indicated. Cell types are labelled and color coded. Note spectrosomes/fusomes in red. **(b,c)** Confocal z-stack images of control **(b)** and *tai^PPxA^* **(c)** germaria 2-days AAE, with anterior positioned to the left. DAPI (blue) marks nuclei, Vasa (green) labels germline cells and α-Spectrin (αSpec; magenta) labels spectrosomes and fusomes. White arrows in *tai^PPxA^*(c) indicate dot spectrosomes. **(c’-d’)** Inverted greyscale of αSpectrin channel for control and *tai^PPxA^* to highlight spectrosomes and fusomes. Red arrows in (**c’**) indicate mature fusosmes that wrap control CB clusters, and are missing in *tai^PPxA^* germaria (**d’**). **(d)** Quantification of dot spectrosome-containing cells per germarium, showing an accumulation of undifferentiated germline cells in *tai^PPxA^* mutants. Sample sizes: control: n=12; *tai^PPxA^*: n=27. Scale bar, 10 μm. **(e)** Diagram of stage 9-10 unfertilized oocyte undergoing border cell (BC) migration. The BC cluster is green, and other cell types are labelled and colored. **(f-g)** Confocal images of control (**f**) and *tai^PPxA^* (**g**) stage 10 egg chambers. *slbo>GFP* (green) labels the BC cluster and some follicle cells, Armadillo (Arm; magenta) labels cell-cell junctions. Scale bar 50 μm. **(h)** Percent migration of cells in stage 10 egg chambers. Sample sizes: control n=27, *tai^PPxA^* n=36. **(g, h)** Statistical significance was assessed using two-tailed unpaired t-tests. Error bars represent SEM. * = P ≤ 0.05; ns, not significant.

When applied to control germaria, anti-αSpec labels 2-3 dot spectrosomes in the anterior germarium, small immature fusomes in the mid region, and mature branched fusomes spanning CB clusters located in the posterior germarium (**Fig. 2b-b’**). By contrast, *tai^PPxA^* germaria contain excess dot spectrosomes (avg. 5.7±1.6 versus avg. of 3.4±0.9 in control germaria, p=2.3e-6, **Fig. 2d**) and a concomitant loss of branched fusomes (**Fig. 2b-b’, c-c’**). This difference in αSpec staining and spectrosome maturation indicates that *tai^PPxA^* likely does not block GSC divisions but rather impairs differentiation of cystoblasts as they move posteriorly away from the niche.

To assess whether *tai^PPxA^* broadly disrupts Tai function, we examined the Tai-dependent process of border cell (BC) migration in the female germline. BC migration occurs during late egg maturation (stage 9-10) and involves a group of ovarian follicle cells that delaminate and migrate between nurse cells to the posterior edge of oocyte (see cartoon, **Fig. 2e**) (rev. in Montell, 2003). This process is blocked or slowed by loss-of-function alleles of *tai* or *EcR*, or local loss of the cytochrome P450 enzyme *shade* which converts the steroid hormone ecdysone to its active 20-hydroxyecdysone form (Bai et al., 2000b; Bhattacharya and Starz-Gaiano, 2024; Domanitskaya et al., 2014b; Hackney et al., 2007). A *slbo-GFP* transgene (*slow border cells*) (Montell et al., 1992) was used to track BC migration in *control* and *tai^PPxA^* egg chambers that reach stage 10, when the BC cluster reaches the immature oocyte (rev. in Montell, 2003). Relatively few *tai^PPxA^* mutant egg chambers progress to stage 10. Those that do show a slight but non-significant reduction in BC migration relative with controls (p=0.084; **Figs. 2f-h**), suggesting that *tai^PPxA^* has a more subtle effect on BC migration than CB differentiation. These data indicate that the Yki-interaction motif in Tai has a more significant role in GSC differentiation than BC migration.

### Loss of the Yki:Tai interaction in ECs leads to germline differentiation defects

Tai protein is highly expressed in anterior ECs but also present at lower levels in other cells in the germarium (Song et al., 2007). To assess whether disruption of the Yki:Tai interaction in subtypes of germarial cells is sufficient to perturb GSC differentiation, we targeted *tai^PPxA^* to groups of cells using a *UAS-tai^PPxA^* transgene (described in Zhang et al., 2015a) and *Gal4* lines that direct expression predominantly in either anterior (*C587-Gal4*) or posterior ECs (*ptc-Gal4*), in cap cells and TF cells (*bab1-Gal4*), in follicle cells (*tj-Gal4*), or in GSCs (*otu-Gal4* and *nanos-Gal4*) (**Fig. 3a-j**). The *bab1>tai^PPxA^* combination was larval lethal, likely due to its additional strong expression in brain and gut (Weaver et al., 2020) and thus could not be analyzed. Analysis with αSpec and Vasa antibodies (**Fig. 3a-j**) detected accumulation of dot spectrosomes in *C587>tai^PPxA^* ovaries that resembles the accumulation of dot spectrosomes in fully *tai^PPxA^* mutant germaria (**Fig. 3b,g**). As noted above, the *C587-Gal4* driver is active in anterior ECs (Diaz-Torres et al., 2021; Mao et al., 2019; Tu et al., 2023; Tu et al., 2026) which directly contact GSCs and exhibit a different pattern of gene expression relative to posterior ECs, including higher levels of *tai* expression (Song et al., 2007; Tu et al., 2021). Notably, expression of *tai^PPxA^* from the posterior EC driver *ptc-Gal4* did not alter dot spectrosomes (**Fig. 3a,f**), perhaps due to lower activity of the *ptc-Gal4* driver in anterior ECs (Weaver et al., 2020). In sum, these data indicate that cystoblast differentiation requires Tai to be competent to bind Yki in a *C587-Gal4*^strong^/*ptc-Gal4*^weak^ subset of ECs.

**Figure 3.**
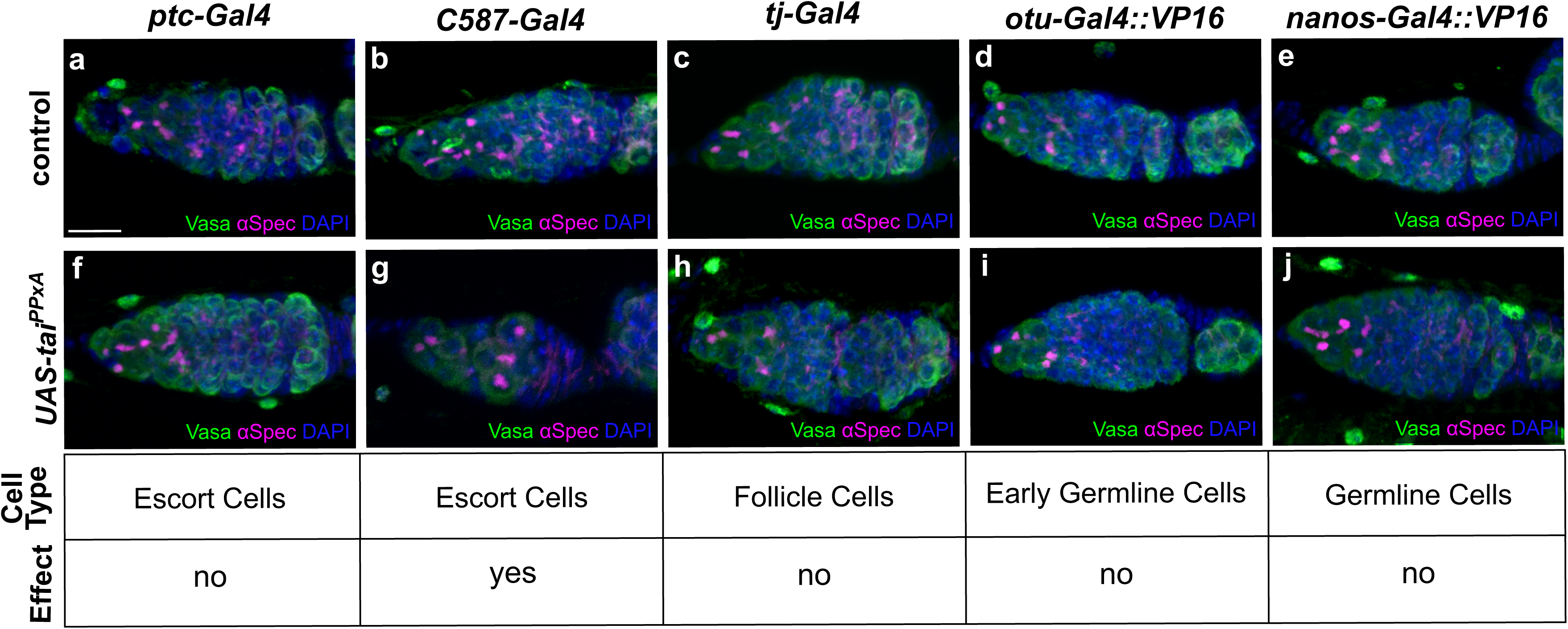
Expression of a *UAS-tai^PPxA^* transgene in ECs causes germline differentiation defects. (**a-j**) Germaria from wildtype females with the indicated Gal 4driver alone (**a-e**) and in combination with *UAS-tai^PPxA^* **(f-j)**. The *C587-Gal4,UAS-tai^PPxA^* combination **(g)** produced small germaria with extra dot spectrosomes and lack of branched fusomes. All images represent a confocal z-stack. Scale bar, 10 *μ*m.

### The Yki:Tai interaction maintains EC morphology in adulthood

Depletion of Yki from anterior ECs shortens their cytoplasmic processes that normally ensheathe germline cysts (Huang et al., 2017), and this altered EC morphology is associated with defects in GSC differentiation (Kirilly et al., 2011; Maimon et al., 2014; Schulz et al., 2002). Therefore, we examined the effects of *tai^PPxA^* expression on EC processes by comparing the effects of *UAS-tai^PPxA^* and wildtype *tai* (*UAS-tai^wt^*) transgenes using the anterior EC driver *C587-Gal4*. Due to the lethality of prolonged overexpression of *UAS-tai^wt^*, we utilized the *tubulin-Gal80^ts^* system (Suster et al., 2004) to restrict *tai* transgene expression to adult ECs. After raising unmated females at the permissive temperature (18°C), we shifted newly eclosed adult females to the restrictive temperature (29°C) for 3 days and then analyzed EC two-dimensional volume (cell bodies+processes) relative to the overall size of the host germaria, which we term ‘relative EC area’. Relative EC area was slightly reduced in 3-day old *tubGal80^ts^;C587-Gal4>tai^wt^* germaria (p=0.018; **Fig. 4b-b’’,4d**) and more strongly reduced in 3-day *old tubGal80^ts^;C587-Gal4>tai^PPxA^* germaria. This effect of *tai^PPxA^* on relative EC area was significant compared to both *control* and *tai^wt^* germaria (p=1.1e-6 and p=2.2e-9 respectively; **Fig. 4c-c’’,4d**). However, no significant difference in spectrosome number was observed in either *tubGal80^ts^;C587-Gal4>tai^wt^*or *tai^PPxA^* backgrounds after 3-days of exposure to the restrictive temperature, likely due to the limited duration of transgene expression.

**Figure 4.**
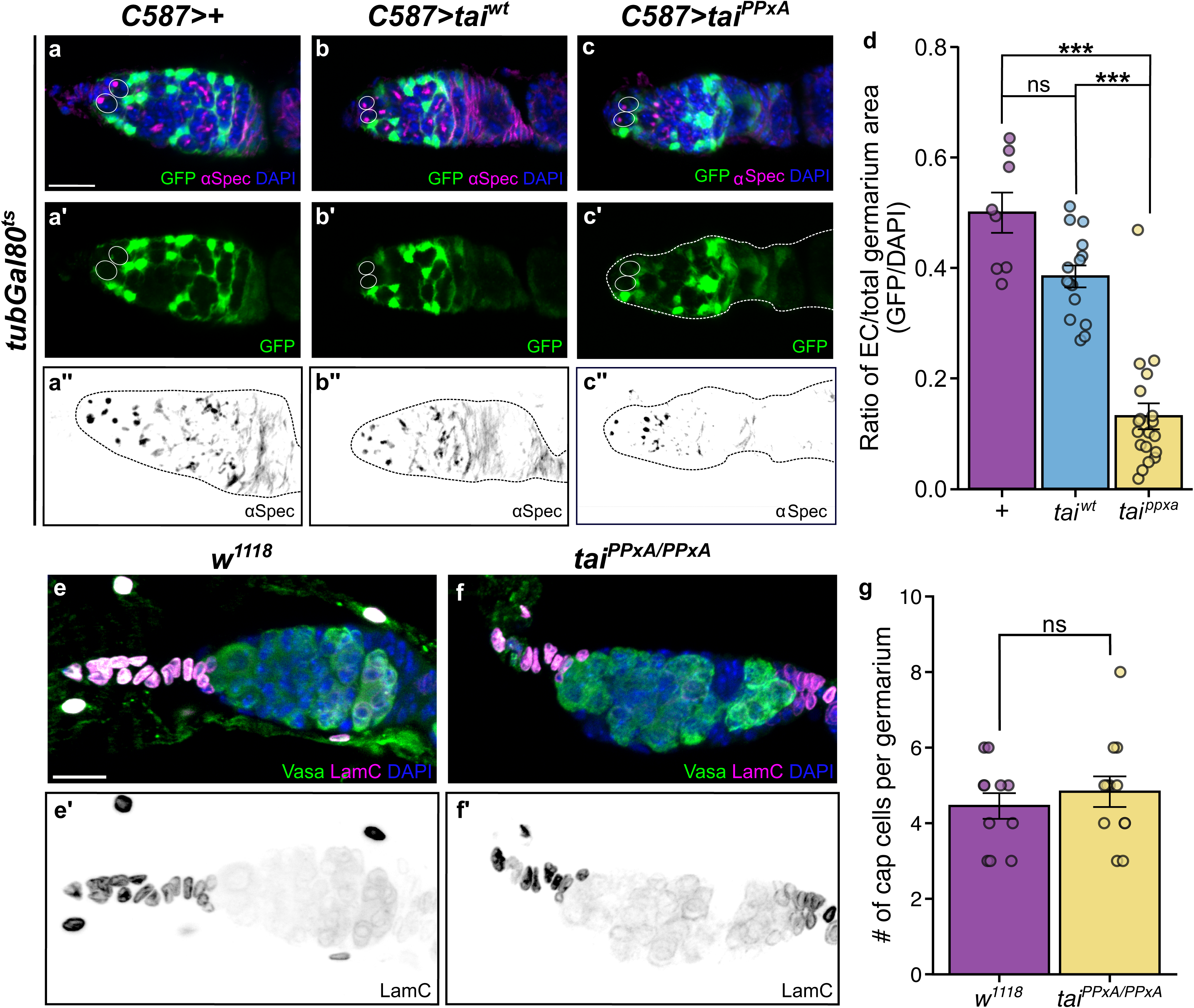
The Yki:Tai interaction maintains EC morphology in adulthood. **(a-c)** Single confocal slices of germaria from females carrying the *C587-Gal4*+*tub-Gal80^ts^*transgene combination alone **(a)**, or in combination with *UAS-tai^wt^***(b)**, or *UAS-tai^PPxA^* **(c)**. Adults were reared at 18° and shifted to 29°C for three days prior to analysis. ECs are labelled GFP (green; *UAS-GFP*), αSpectrin (magenta) labels spectrosomes and fusomes, and DAPI marks nuclei. Circles indicate GSCs. **(a’-c’)** Single slices of GFP channel marking only ECs in *C587,tubGal80^ts^>+* (control), *C587,tubGal80^ts^>tai^wt^*, and *C587,tubGal80^ts^>tai^PPxA^* germaria. **(a’’-c’’)** Confocal z-stacks of GFP channel marking only ECs in *C587,tubGal80^ts^>+* (control), *C587,tubGal80^ts^>tai^wt^*, and *C587,tubGal80^ts^>tai^PPxA^* germaria. **(a’’’-c’’’)** Confocal z-stacks of inverted greyscale of αSpectrin signal for the same genotypes as in **a-a’’**, **b-b’’**, and **c-c’’** highlighting spectrosomes and fusomes. **(d)** Quantification of EC area showing that three-day induction of *UAS-tai^PPxA^* germaria significantly reduces the EC/germarium area ratio compared to control and *UAS-tai^wt^*. Samples sizes: control (n = 8), *UAS-tai*^wt^ (n = 15), *UAS-tai^PPxA^* (n = 19). **(e-f)** Confocal z-stacks of germaria from control **(e)** and *tai^PPxA^***(f)** females labelled with LaminC (magenta) to detect cap and terminal filament cells, Vasa (green) to mark germline, and DAPI to mark nuclei. **(e’-f’)** Inverted grayscale of LamC channel for control (**e’**) and *tai^PPxA^* (**f’**) females. **(g)** Quantification of the number of cap cells (smaller that TF cells) in control (*w^1118^*) and *tai^PPxA^* germaria. Samples sizes: control (n=11), *tai^PPxA^* (n=12). Statistics were calculated using Welch’s unpaired t-tests with Bonferroni correction. Scale bars,10 µm. Error bars represent *SEM*. * = P ≤ 0.05, ** = P ≤ 0.005, *** = P ≤ 0.0005; ns, not significant.

Elevating cap cell numbers above the 5-7 average per niche can produce defects in GSC differentiation due to their role as the primary source of Dpp ligands (Song et al., 2007; Song et al., 2004; Ward et al., 2006; Xie and Spradling, 1998b, 2000) . Although Yki-depletion does not significantly alter cap cell numbers (Huang et al., 2017), elevating Tai expression does expand cap cell number (König et al., 2011). To better assess whether the observed increase in dot spectrosome-containing cells in *tai^PPxA^* germaria could be due to expansion of the cap cell population, we used the cap cell marker Lamin C (LamC) to assess cap cell number and localization. We found that control germaria contained an average of 4.5±1.1 cap cells (**Fig. 4e-e’, 4g**) while *tai^PPxA^* germaria contained an average of 4.8±1.4 cap cells (p=0.49 between genotypes; **Fig. 4f-f’, 4g**). This rather mild difference in cap cell number suggests that the effect of *tai^PPxA^* on GSC differentiation is primarily due to an effect on *C587-Gal4* positive anterior ECs.

### The Yki:Tai interaction limits BMP signaling to the anterior niche

The accumulation of dot spectrosome-containing cells in *tai^PPxA^* germaria raises the possibility that the Tai PPxY motifs support CB differentiation by blocking prolonged BMP exposure. As a preliminary test of BMP signaling, we dissected control and *tai^PPxA^* whole ovaries and performed RT-qPCR analysis on key BMP pathway components (**Fig. S2**), including *dpp* itself, the extracellular glypican *dally* that facilities diffusion of Dpp through the apical ECM (Hayashi et al., 2009; Su et al., 2018; Tsuda et al., 1999), the piRNA-interacting factor *piwi* that represses *dpp* mRNA levels in ECs (Ishizu et al., 2012; Ma et al., 2014), the *upd3* Jak/STAT ligand that promotes *dpp* transcription in cap cells (Wang et al., 2008), as well as the Yki-target gene *expanded* (Hamaratoglu et al., 2006). *tai^PPxA^* did not significantly alter levels of these mRNAs relative to control ovaries, which could either indicate that the PPxY motifs are not involved in expression of Dpp signaling components or that such a role is limited to a small subset of cells (e.g. anterior ECs). To bypass this practical limitation of cellular heterogeneity, we visualized the extracellular Dpp gradient directly using a viable CRISPR-tagged *dpp* allele (*dpp^mCherry^*; BDSC #86273) and the Dpp-responsive *Daughters-against-dpp* (*Dad*)-*lacZ* reporter line to detect the pattern of Dpp signaling in germarium cells along the anterior-posterior (A-P) axis of the germarium. *Dad* is a direct transcriptional target of BMP signaling and only expressed in GSCs in response to short-range BMP ligands secreted by adjacent cap cells (Tsuneizumi et al., 1997; Xie and Spradling, 1998a). As expected, control germaria contain extracellular Dpp^mCherry^ concentrated around cap cells and immediately adjacent to GSCs in the niche (asterisks in **Fig. 5a-b**). *tai^PPxA^* expands this extracellular Dpp^Cherry^ gradient well beyond niche-associated GSCs and into more posterior regions of the germarium (asterisks in **Fig. 5c-d**). Quantification of extracellular Dpp^mCherry^ intensity along the A-P axis indicates that the Dpp^mCherry^ peak is higher in *tai^PPxA^*germaria than control germaria, leading to a posterior extension of the resulting gradient with *tai^PPxA^* germaria exhibiting significantly higher mCherry intensity in four of five distance ‘bins’ spanning the germarium (**Fig. 5e**). In parallel, *tai^PPxA^* homozygous germaria contain an increase in number of *Dad-lacZ* positive cells (i.e., βGal-positive) from 2-3 in control germaria to ∼5 in *tai^PPxA^* germaria (**Fig. 5f-g**). In sum, these data provide evidence that uncoupling the Yki:Tai interaction ectopically expands the gradient of extracellular Dpp in the germarium, which correlates with delayed differentiation of CBs and loss of mature oocytes, and raises the possibility that the Yki:Tai interaction acts upstream of Dpp gradients in other tissues as well .

**Figure 5.**
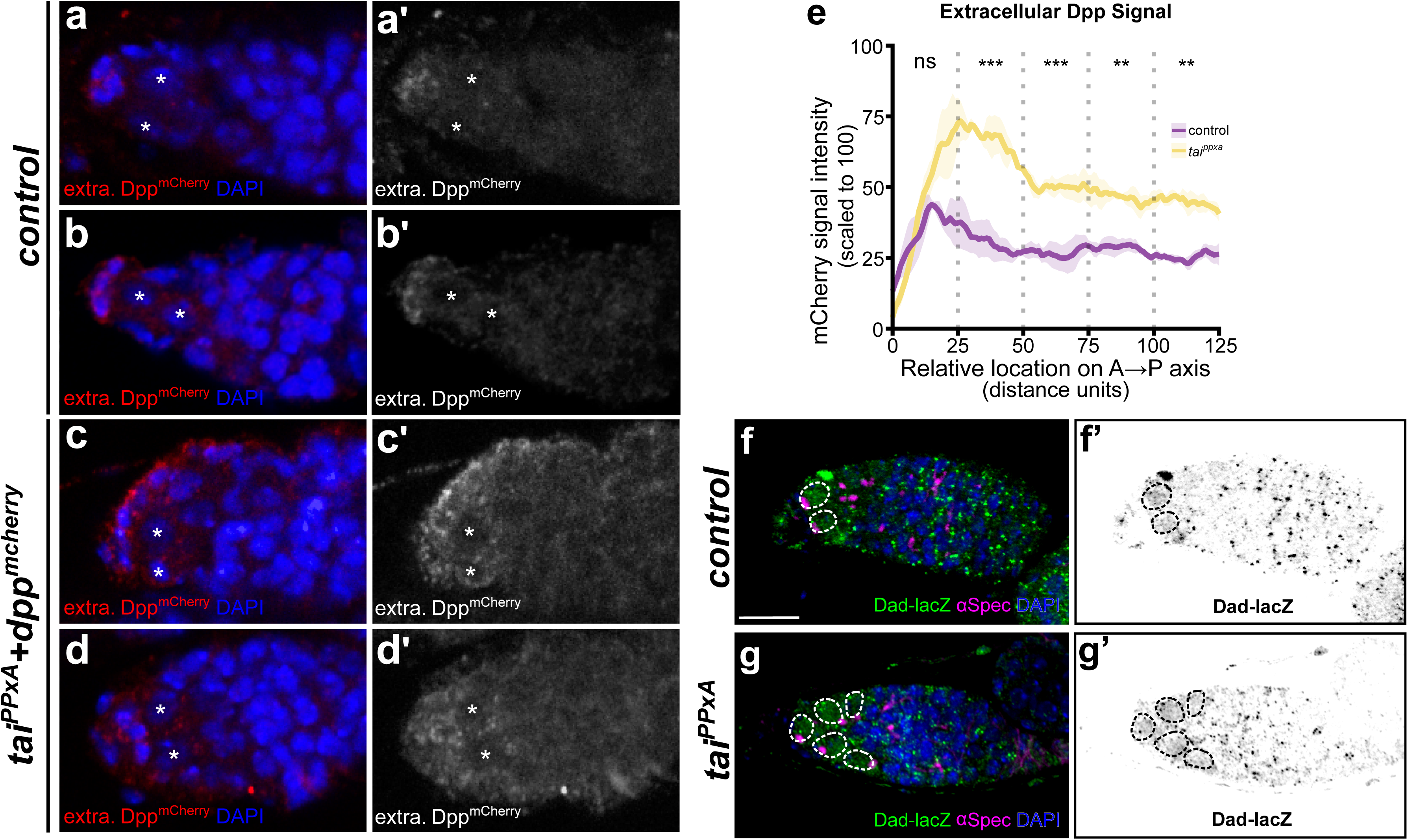
*tai^PPxA^* expands the extracellular Dpp gradient beyond the GSC niche. **(a-d)** Confocal z-stack images of two control (*w^1118^*) (**a-b**) and two *tai^PPxA^* (**c-d**) germaria stained with anti-RFP to detect Dpp^mCherry^ (red) and DAPI (blue) to mark nuclei. GCSs are indicated by asterisks. Samples were fixed but not permeabilized and processed according to (Witte et al., 2021) to selectively detect extracellular Dpp^mCherry^. (**e**) Quantitation of Dpp^mCherry^ signal intensity along the anterior to posterior (A→P) axis of *dpp^mCherry^* (magenta plot) and *tai^PPxA^,dpp^mCherry^* (yellow plot) germaria, with the X-axis “0-value” corresponding to the anterior tip. The region of the germarium imaged in (**a-d**) was split into quintiles (dotted grey lines) that were tested for pairwise significance using a standard T-test. * = P ≤ 0.05, ** = P ≤ 0.005, *** = P ≤ 0.0005; ns, not significant. **(f-g)**. Representative confocal z-stacks of germaria from control (**f**) and *tai^PPxA^* mutants (**g**) expressing β-Galactosidase (β-Gal; green) under the control of the Dad promoter. α-Spectrin (magenta) labels spectrosomes and fusomes, and DAPI (blue) marks nuclei. **(f’-g’)** Inverted greyscale of anti-β-Gal signal for control (**f’**) and *tai^PPxA^* (**g’**). Dotted circles highlight *Dad-lacZ* expressing cells. Scale bar,10 *μ*m.

## Discussion

Stem cell maintenance requires integration of extrinsic signals from the surrounding niche that balance self-renewal and differentiation. In the *Drosophila* germarium, BMP signaling is a central regulator of GSC identity, supporting GSC self-renewal by repressing differentiation and ensuring stem cell homeostasis. Accordingly, BMP signaling is subject to regulation by multiple niche-associated factors, including Hedgehog (Liu et al., 2015), Jak/STAT (Wang et al., 2008), Wg/Wnt (Hamada-Kawaguchi et al., 2014), EGFR (Banisch et al., 2017), Notch (Song et al., 2007), Lsd1 (Eliazer et al., 2011), Piwi (Jin et al., 2013), Hippo (Huang et al., 2017), and Ecdysone signaling (König et al., 2011). Our findings identify a point of convergence between Hippo and Ecdysone signaling through the interaction of their respective transcriptional coactivators Yki and Tai, mediated by two WW domains in Yki and two PPxY motifs in Tai. We find that disruption of the Yki:Tai interaction is associated with altered EC morphology, impaired GSC differentiation, and ectopic BMP signaling, supporting a model in which Yki:Tai crosstalk in ECs contributes to the spatial restriction of niche signals within the germarium.

EC projections that extend between developing germline cysts help restrict exposure of CBs to niche-derived extracellular BMP ligands as CBs move posteriorly through the germarium (Kirilly et al., 2011; Morris and Spradling, 2011). Previous studies demonstrated that disruption of either Yki or Tai shortens these EC projections and impairs germline differentiation (Huang et al., 2017; König et al., 2011). Our data provide mechanistic insight into these roles by demonstrating that they are based on a pair of WW:PPxY interactions between Yki and Tai proteins. EC-specific expression of *tai^PPxA^* resulted in shortened EC processes that fail to project fully around CBs in the germarium. This morphological phenotype is accompanied by an accumulation of dot spectrosome-containing CB cells, which is consistent with impaired progression of GSC daughter-cell differentiation. In addition to these EC morphological changes, *tai^PPxA^* expands the spatial distribution range of BMP signaling. Levels of extracellular Dpp are elevated and extend more posteriorly in *tai^PPxA^* germaria, which is correlated with an accumulation of “dot spectrosomes” and expression of the BMP-induced gene *Dad* in posterior cells. Overall, these data indicate that disrupting the Yki:Tai interaction represses the level and range of BMP signaling in a manner that phenocopies loss of either Tai or Yki, which in turn suggests that the Yki:Tai complex is a key functional unit in ECs. Importantly, *tai^PPxA^* retains ability to coactivate EcR-dependent processes such as BC migration and GSC proliferation (Ameku and Niwa, 2016a; Bai et al., 2000a; Domanitskaya et al., 2014a; Jung et al., 2026) and has no effect on cap cell number, which is increased in germaria expressing *tai* RNAi transgene (König et al., 2011). These phenotypic differences between *tai* RNAi and *tai^PPxA^* supoorts that Tai^PPxA^ remains competent to function as an EcR coactivator. Whether the reciprocal role of Tai role in co-activating transcription of Hippo target genes involved in wound healing and tissue renewal (Wang et al., 2016; Zhang et al., 2015a) requires a Tai:EcR interaction is not known, but remains a possibility based on our previous data that Tai can simultaneously associate with Yki and EcR (Zhang et al., 2015a).

The transcriptional mechanism by which the Yki:Tai complex controls Dpp signaling remains unresolved. The ability of *tai^PPxA^*to replicate effects of Yki or Tai RNAi in ECs implies that one effect of *tai^PPxA^*is to block cooperative induction of shared targets, as occurs when Tai^PPxA^ is co-overexpressed with Yki in larval wing cells (Wang et al., 2016; Zhang et al., 2015a). Canonically, Yki regulates transcription through its DNA-binding partner Scalloped (Sd). The lack of a “dot spectrosome” defect in germaria with Sd-depleted ECs has led to speculation that Yki acts via a non-canonical mechanism in these cells (Huang et al., 2017). However, as Sd can recruit both activators and repressors to Hippo target genes (Guo et al., 2013; Koontz et al., 2013), Sd depletion in ECs may produce deficits in activation and repression that counterbalance each other on relevant target genes. Resolving these questions will require identification of direct Yki:Tai target genes in ECs. When analyzed at the level of whole *wt* and *tai^PPxA^* ovaries, *tai^PPxA^* has only weak effects levels of mRNAs encoding Dpp and the Dally glypican, which is an extracellular glypican that binds Dpp and facilitates extracellular across multiple tissues and presents it to the Dpp receptor Tkv on GSCs (Belenkaya et al., 2004; Su et al., 2018). While there are multiple Dpp regulators that could be tested at the level of whole ovaries, these initial data suggest either that *tai^PPxA^* enhances Dpp signaling through an unknown intermediate, or that its effect on gene expression is restricted to a small subset of cells that will need to be identified by other means (e.g., FISH or a survey of candidate enhancer trap lines). Interestingly, Tai controls levels of the Dally paralog Dally-like-protein (Dlp) (Schweibenz et al., 2024) in larval wing disc cells, suggesting that *tai* may regulates some aspect of glypican production in ECs (e.g., secretion, recycling, or glycosylation). Further work is required to understand how Yki:Tai controls the germarial Dpp gradient, and what role Yki:Tai and Tai:EcR interactions play in coordinating expression of ecdysone and Hippo target genes that are crucial for maintenance of the germline stem cell niche.

## Materials and Methods

### Drosophila culture

Fly stocks were maintained under standard culture conditions at 25°C in humidity-controlled incubators with 12-hour light/dark cycles. All crosses were maintained on standard molasses food supplemented with yeast. Unless otherwise indicated, all ovary dissections were from unmated females 2-3 days after adult eclosion maintained on standard food with yeast paste (1:1 mixture of water and dry yeast flakes). The following strains were used: *C587-Gal4* (BDSC #67747), *ptc-Gal4* (BDSC #2017), *tj-Gal4* (BDSC #50105), *bab-Gal4* (BDSC #6803), *out-Gal4::VP16* (BDSC #58424), *nanos-Gal4::VP16* (BDSC #4937), *Dad-lacZ* (BDSC #10305), *UAS-tai* (Zhang et al., 2015b), *UAS-tai^PPxA^* (Zhang et al., 2015b), *tubGal80ts, UAS-GFP* (BDSC #67072), *dpp^mCherry^* (BDSC #86273), *tai^PPxA^* (this study).

### CRISPR editing of endogenous *tai*

CRISPR editing was performed by Rainbow Transgenics, Inc and verified by direct sequencing. Briefly, a sgRNA targeting a PAM site between the two PPxY motifs of *tai* was co-injected with a homology directed repair plasmid into *vasa-Cas9* embryos. The HDR plasmid contained PPxY-to-A edits, an edited PAM site to prevent additional targeting events, and a *3xP3>GFP* cassette flanked by *loxP* sites, all of which were inserted into a flanking intron. Successful targeting events were identified by visual scoring for GFP and then crossed to Cre recombinase lines (BDSC #766). Lines were confirmed by sequencing the *tai* locus and then backcrossed 8 times to a wildtype *iso2:iso3* background and balanced.

### Ovary size determination

Control and mutant unmated females were collected and incubated with yeast paste 3 days before dissection at 3, 7, or 14 days after adult eclosion. Whole ovaries were dissected in 1X phosphate buffered saline (PBS) and imaged in a 9-well PYREX plate on a Nikon SMZ800N at 3X magnification. The area of each ovary was determined on Adobe photoshop using the lasso tool.

### Immunofluorescence and microscopy

Immunofluorescence and microscopy were performed using standard methods. Ovaries were dissected in 1X PBS, fixed for 20 minutes in 4% paraformaldehyde (PFA) at room temperature, rinsed 3X in 1X PBS, and then permeabilized with 1X PBS and 0.3% Triton X-100 (0.3% PBST). Ovaries were then blocked with 10% normal goat serum (NGS) and incubated in diluted primary antibody containing 10% NGS and 1X PBS with 0.1% Triton X-100 (0.1% PBST). Samples were incubated in primary antibody solution at 4°C overnight, then washed 5X with 0.1% PBST, incubated with DAPI (1:1000) for 10 minutes, washed 3X with 0.1% PBST, and then incubated for 1 hr at RT with secondary antibody solution diluted with 10% NGS and 0.1% PBST. Ovaries were then washed 5X times with 0.1% PBST and incubated in n-propyl gallate (NPG, 4% w/v in glycerol) overnight. Detergents were omitted from experiments to detect extracellular Dpp^mCherry^ {as in \Witte, 2021 #4579}. Ovaries were mounted in NPG on glass coverslips and then imaged with a Nikon A1R confocal system using 40X objectives in oil. Images were processed with Fiji and displayed with Inkscape. The following antibodies were used: Developmental Studies Hybridoma Bank mouse anti-Spectrin (1:200), rat anti-Vasa (1:10), and Invitrogen rabbit anti-β-galactosidase (1:500), rabbit anti-GFP (1:1000), goat anti-mouse Cy3 (1:50), goat anti-mouse Cy5 (1:50), goat anti-rabbit Cy3, DAPI (1:1000). Other antibodies used: Goat anti-rat 488 (1:500; gift of D. Lerit).

### Disruption of Yki:Tai interaction in soma

To specifically disrupt the Yki:Tai interaction in stomatic cells of the germarium, *UAS-tai* or *UAS-tai^PPxA^* males were crossed to *C587-Gal4;tgGal80ts,UAS-GFP/TM2* females at the permissive temperature (18°). Hatched flies were then collected and transferred to the restrictive temperature (29°) for 3 days before dissection. Controls were treated the same.

### Quantitative Reverse Transcription PCR (RT-qPCR)

We dissected whole ovaries from unmated females 2-3 days AAE into TriZol (Invtirogen; 15596018), and RNA was extracted using the Qiagen RNeasy kit (Qiagen; 74106) according to the manufacturer’s instructions. cDNA conversion was performed using a SuperScript-III RT kit (Thermo Fisher Scientific; 18080093) according to the manufacturer’s instruction using 500 ng of extracted RNA. qPCR was performed in technical triplicate using a SYBR Green Master Mix (Qiagen, 204143) on a Flex real-time PCR system (Applied Biosystems QuantStudio6). qPCR reactions were carried out for 30 mins at 50 degrees, followed by 10 min at 95 degrees, then 40 cycles of two-step PCR for 15s at 95 degrees, 1 min at 60 degrees. RNA levels were normalized to rp49 and relative concentration was determined using the 2−ΔΔCt method (Livak and Schmittgen, 2001).

The following primers were used:

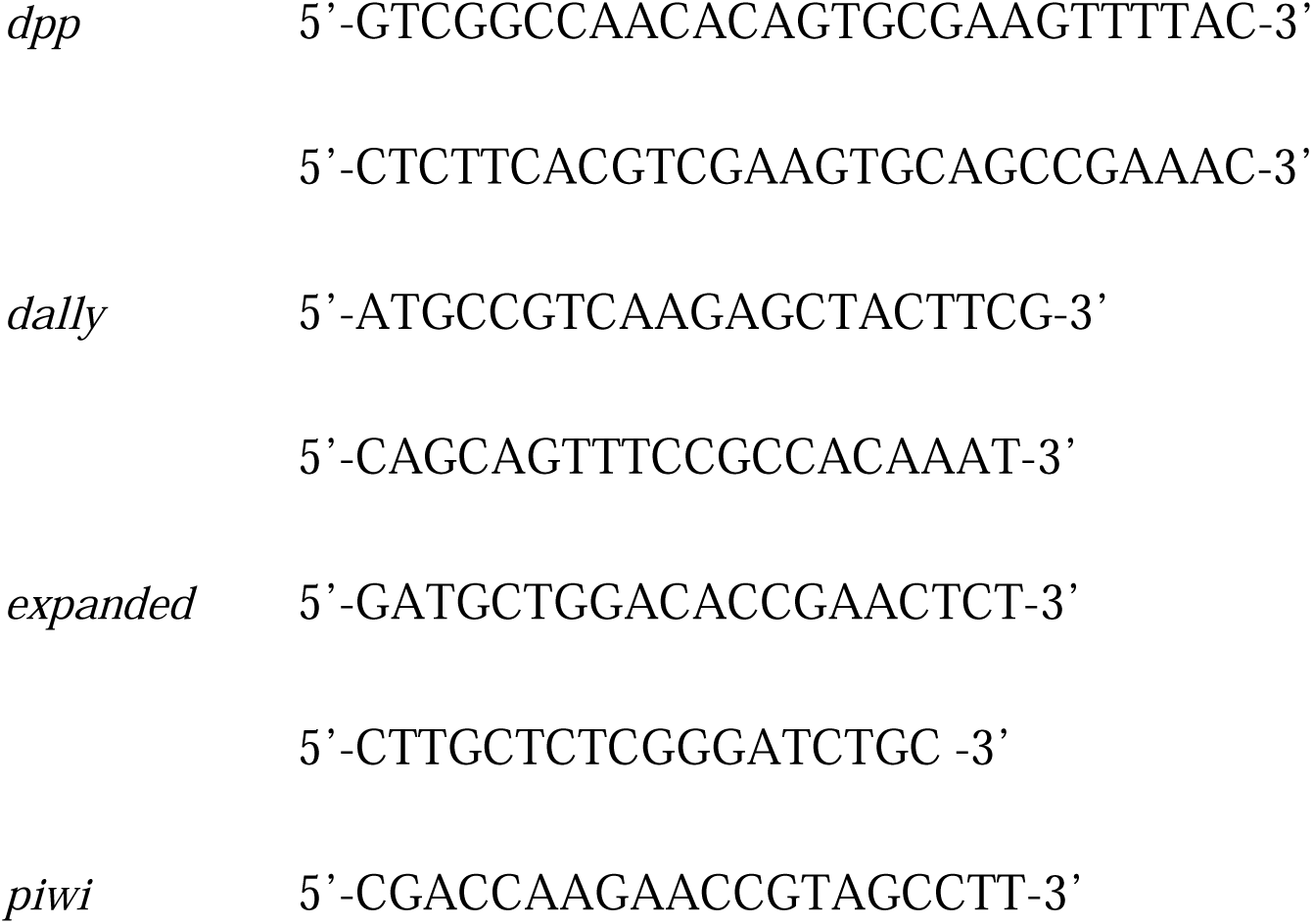
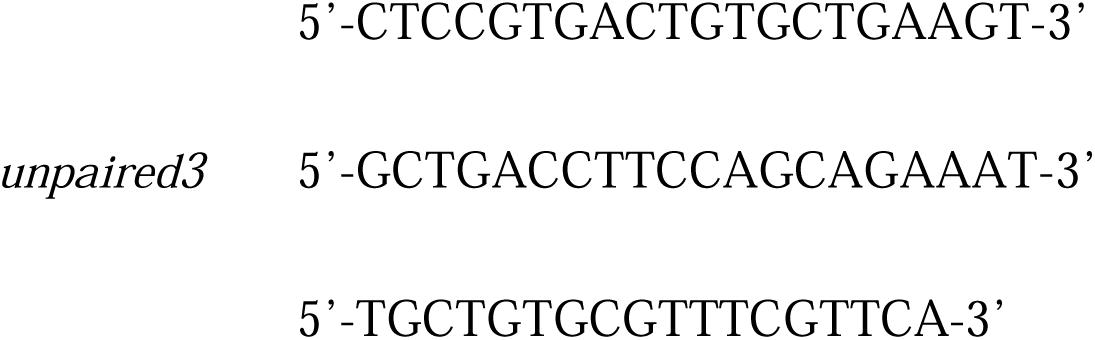

### Analyses and statistics

To determine dot spectrosome count, Z-stacks of αSpectrin stained ovaries were taken every 1µm, spherical appearing spectrosomes were counted that were not present in over three slices (3µm), to account for branched fusomes presenting as spherical spectrosomes. To determine escort cell area, germarium images were first cropped to the end of region III (depicted in **Fig. 2a** cartoon). Using Fiji, the GFP channel from *C587>GFP* germaria was thresholded, with a lower threshold value that included all real signal and excluded background. The area of the threshold was measured for each germarium. To account for changes in germarium size, the DAPI channel was thresholded and measured, to find the area of the entire germarium. Normalized EC area was calculated by GFP area/germarium area. The same threshold values were used for all genotypes. No samples were excluded from analysis. Other statistical tests are described in each figure legend and calculated using R.

The extracellular distribution of Dpp^mCherry^ was analyzed with the Fiji Plot Profile tool to determine the anterior-to-posterior intensity plot within a fixed-size rectangle anchored on the first terminal filament nuclei at the anterior tip of each germarium. Intensity values were assigned to 25 distance unit bins, and values within each bin were averaged for each germarium. Data were analyzed using a linear mixed-effects model (lme4) with mean intensity as the response, genotype, distance bin, and their interaction as fixed effects, and germarium as a random intercept. Significance of fixed effects was assessed by Type III ANOVA with Satterthwaite’s approximation for degrees of freedom. Estimated marginal means were calculated for each genotype at each distance bin, and genotypes were compared within each bin using pairwise contrasts with Kenward-Roger degrees of freedom. P-values were adjusted for multiple comparisons across the five bins using the Holm method. No samples were excluded from analysis.

## Author contributions

This study was conceptualized by CCW, VCP, and KHM. Experiments were performed by CCW and VCP. Data analysis performed by CCW and VCP. CRISPR mutant flies were designed by SV and KHM. AlphaFold analysis performed by TB. The manuscript was written by CCW with contributions from all authors.

## Acknowledgments

This work was supported by the Emory University Integrated Cellular Imaging Core (RRID:SCR_023534). The content is solely the responsibility of the authors and does not necessarily reflect the official views of the National Institute of Health. We thank D. Lerit, DSHB, and BDSC for reagents and fly stocks. We also thank members of the Moberg and Lerit laboratories for helpful discussions, and members of CCW’s honors thesis committee, L. Reider and D. Gorkin.

---

**Supplemental Figure 1.** Kaplan-Meier survival curve comparison of lifespan data from control *w^1118^* (purple) and *tai^PPxA^* (yellow) adult male flies. Newly eclosed male flies of each genotype were collected on the same day and the number of dead flies per vial was recorded daily. Flies were maintained on standard food and transferred to fresh vials every other day. Median lifespan is indicated by black dashed line. Shaded regions indicated 95% confidence intervals, n=30 for each genotype, and p-value was derived from log rank test.

**Supplemental Figure 2.** RT-qPCR comparison of relativelevels of the indicated mRNAs in control (*w^1118^*) (magenta) versus *tai^PPxA^* (yellow) ovaries. Total mRNA harvested from dissected ovaries was used for this analysis. Values for each mRNA were normalized to “1” for the *w^1118^* controls. Analysis was conducted in biological triplicates. Errors bars represent SEM. None of the pairwise comparisons are significant using a standard T-test.

