## Supplementary figures and images for "An interaction between Yorkie and Taiman transcriptional coactivators regulates niche-to-stem cell BMP signaling in the female *Drosophila* germline"

### Supplemental Fig 1

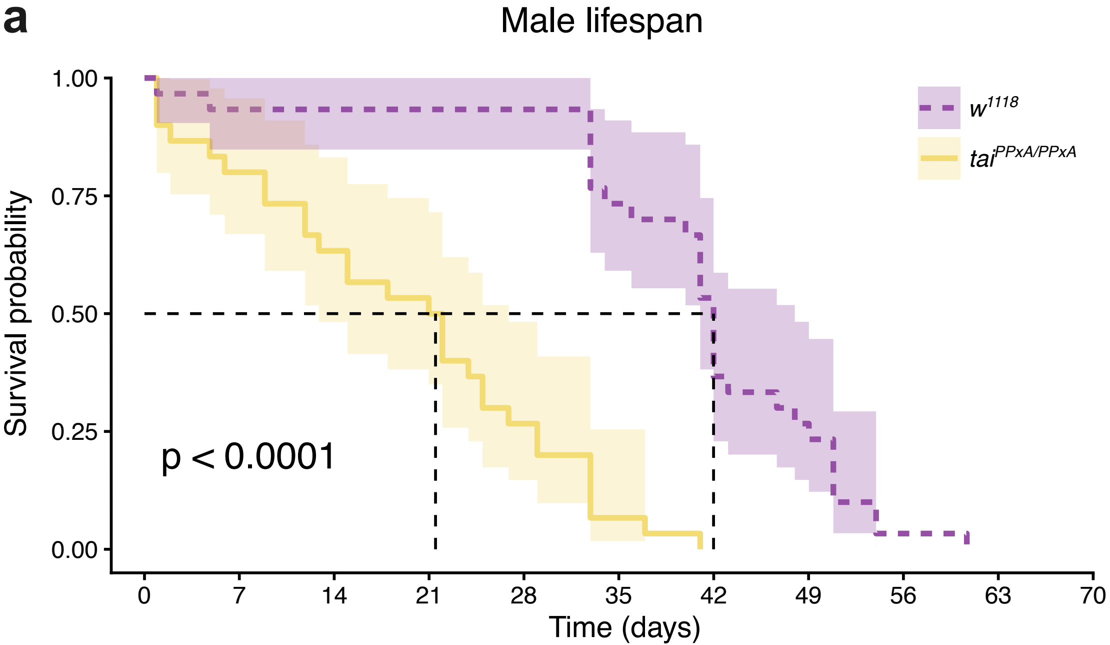

### Supplemental Fig 2

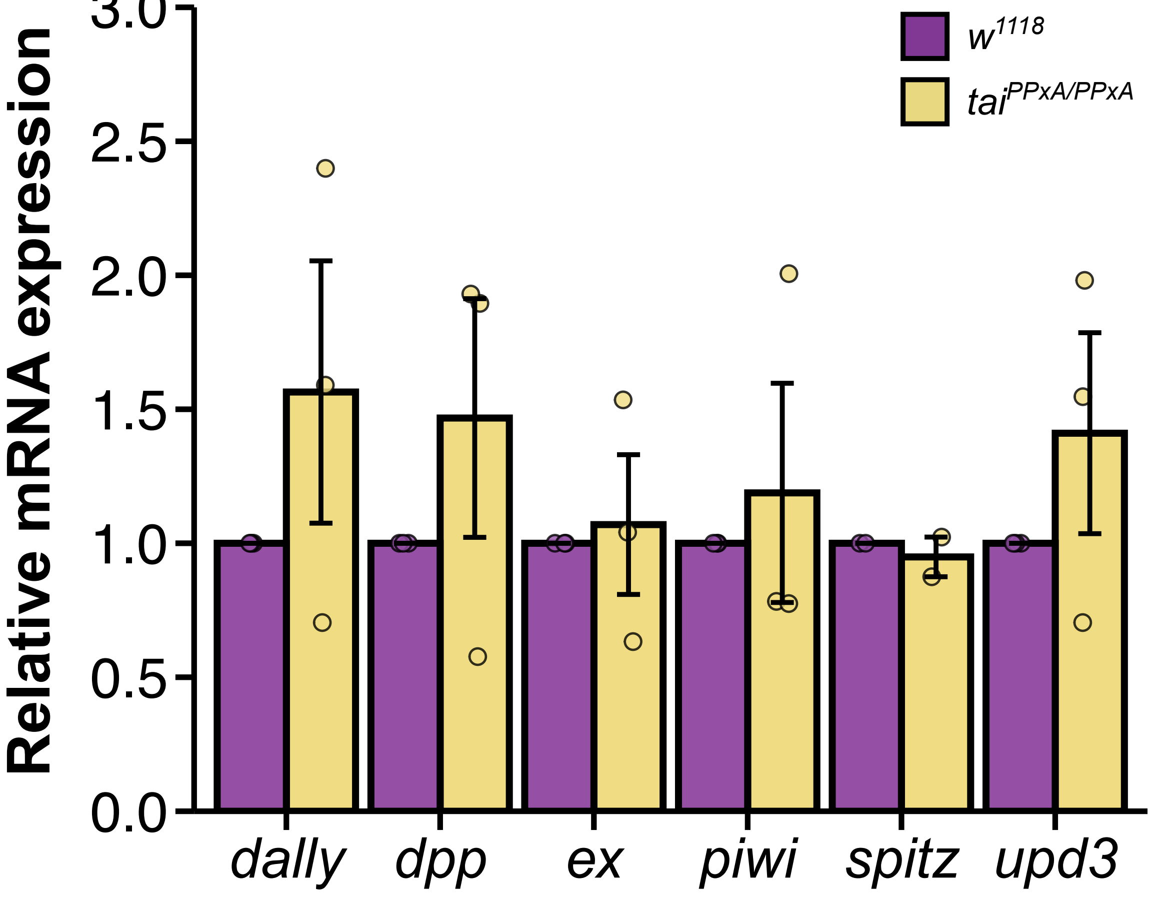
